# Venus-MAXWELL: Efficient Learning of Protein-Mutation Stability Landscapes using Protein Language Models

**DOI:** 10.1101/2025.05.30.656964

**Authors:** Yuanxi Yu, Fan Jiang, Xinzhu Ma, Liang Zhang, Bozitao Zhong, Wanli Ouyang, Guisheng Fan, Huiqun Yu, Liang Hong, Mingchen Li

## Abstract

In-silico prediction of protein mutant stability, measured by the difference in Gibbs free energy change (ΔΔ*G*), is fundamental for protein engineering. Current sequence-to-label methods typically employ the two-stage pipeline: (*i*) encoding mutant sequences using neural networks (*e*.*g*., transformers), followed by (*ii*) the ΔΔ*G* regression from the latent representations. Although these methods have demonstrated promising performance, their dependence on specialized neural network encoders significantly increases the complexity. Additionally, the requirement to *individually compute latent representations for each mutant site* negatively impacts *computational efficiency* and poses the risk of *overfitting*. This work proposes the Venus-Maxwell framework, which reformulates mutation ΔΔ*G* prediction as a sequence-to-landscape task. In Venus-Maxwell, mutations of a protein and their corresponding ΔΔ*G* values are organized into a landscape matrix, allowing our framework to learn the ΔΔ*G* landscape of a protein with a single forward and backward pass during training. Besides, to facilitate future works, we also curated a large-scale ΔΔ*G* dataset with strict controls on data leakage and redundancy to ensure robust evaluation. Venus-Maxwell is compatible with multiple protein language models and enables these models for accurate and efficient ΔΔ*G* prediction. For example, when integrated with the ESM-IF, Venus-Maxwell achieves higher accuracy than ThermoMPNN with 10× faster in inference speed (despite having 50× more parameters than ThermoMPNN). The training codes, model weights, and datasets are publicly available at https://github.com/ai4protein/Venus-MAXWELL.

## 1 Introduction

Protein stability, determined by the change in Gibbs free energy (Δ*G*) of protein sequences between the native and unfolded states, plays a critical role in various applications, ranging from therapeutic protein design to industrial enzyme engineering [1, 2, 3, 4, 5, 6]. In protein engineering, accurately predicting mutation-induced stability changes (ΔΔ*G*) is critical, and current methods for assessing stability across large mutant libraries remain prohibitively expensive and resource-intensive, creating a pressing need for efficient computational solutions. Consequently, developing accurate and efficient computational tools for protein stability prediction is a longstanding focus of research, spanning physics-based and statistical methods [7, 8, 9] to machine learning approaches [10, 11, 12, 13, 14].

Deep learning, exemplified by AlphaFold2 [15], has revolutionized protein-related tasks [16, 17, 18, 19, 20, 21] with protein language models (PLMs), emerging as powerful tools for capturing co-evolutionary patterns from vast sequence and structural datasets [22, 23, 24, 25, 26]. These models have shown powerful capabilities in wide-range downstream tasks, including stability prediction. Recent methods, such as Stability Oracle [13] and ThermoMPNN [14], leverage PLMs to achieve impressive performance in ΔΔ*G* estimation. However, their reliance on complex architectures and massive computational requirements leads to prohibitively high training/inference costs, fundamentally limiting their scalability for large-scale protein engineering applications. These methods can be broadly categorized as sequence-to-label approaches. Specifically, these methodologies involve a two-stage process: first, a PLM or a similar encoder model transforms a mutant sequence into a latent representation; subsequently, a regression head predicts the ΔΔ*G* value from the representation.

The sequence-to-label paradigm, however, presents three main challenges: (*i*) **Computational Inefficiency:** This design requires computing a distinct latent vector for each mutant site (as shown in Figure 1 (A)), which is computationally expensive and heavily dependent on the encoder’s design; (*ii*) **Knowledge Misutilization:** The regression head is often trained from scratch, failing to fully harness the rich evolutionary patterns PLMs learn during pre-training, which are implicitly captured in the model’s output likelihoods or logits. (*iii*) **Architectural Rigidity:** The reliance on a specific regression head design limits versatility and complicates the integration of different PLM types. These limitations highlight the need for a generalizable and efficient approach for stability prediction.

**Figure 1.**
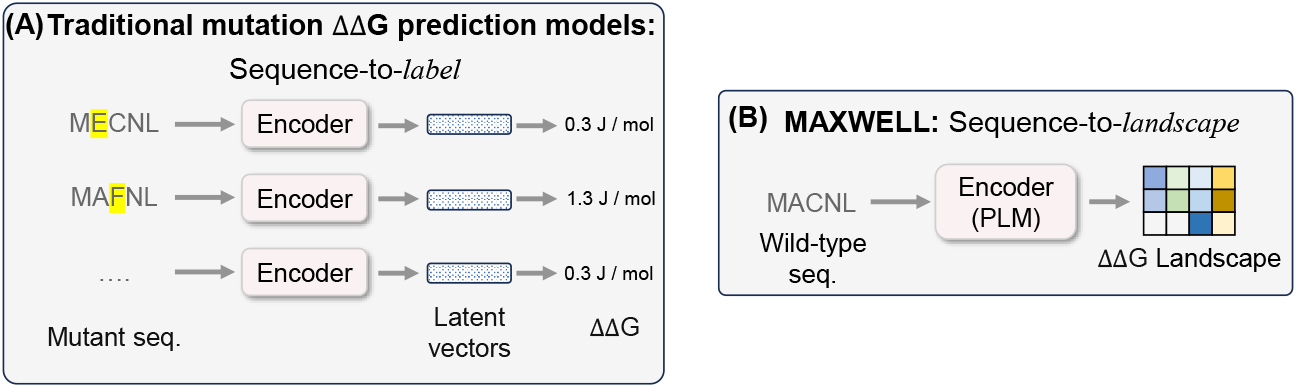
Comparison of Venus-Maxwell and sequence-to-label models. *(A):* The sequence-to-label models compute a latent vector for each mutant site. *(B):* The Venus-Maxwell outputs a landscape in one forward pass.

In this work, we introduce Venus-Maxwell (**Ma**tri**x**-**w**is**e l**andscape **l**earning), an efficient sequence-to-landscape framework that reframes ΔΔ*G* prediction. Venus-Maxwell is not merely a faster model but a new fine-tuning paradigm that directly addresses the limitations of sequence-to-label approaches.

First, Venus-Maxwell maximizes the utilization of pre-trained PLM knowledge. Our key insight is to formulate the framework as a differentiable, matrix-based version of the PLM’s native zero-shot scoring function. Many PLMs, pre-trained on millions of sequences, can estimate the fitness like ΔΔ*G* of mutant in a “zero-shot” manner by calculating the log-likelihood difference between mutant and wild-type sequences[27, 28]. Venus-Maxwell is designed to inherit this powerful capability. Critically, its “initial performance” (before any fine-tuning) is mathematically equivalent to the underlying PLM’s zero-shot prediction performance. Therefore, our fine-tuning process does not learn from scratch but starts from a significantly advanced foundation, directly enhancing the rich evolutionary information already captured by the PLM.

Second, Venus-Maxwell introduces a novel matrix-based fine-tuning methodology. Instead of adding a task-specific regression head, our method directly fine-tunes the PLM’s output logits in a matrix-wise fashion (as depicted in Figure 1 (B)), elegantly adapting the PLM for ΔΔ*G* estimation. This paradigm is highly versatile, serving as a universal fine-tuning framework compatible with diverse PLMs, including masked language models and inverse folding models.

Third, as a direct benefit of this architecture, Venus-Maxwell is exceptionally computationally efficient. To predict all single-site mutations for a protein of 200 amino acids (3800 mutants), a sequence-to-label model must execute 3800 separate computations. In stark contrast, Venus-Maxwell accomplishes all predictions in one computational pass using the wild-type sequence, enabling high-throughput ΔΔ*G* prediction at minimal cost. This capability is particularly important for protein engineering workflows such as early-stage directed evolution, which often begin by comprehensively screening single-site mutations to identify stabilizing candidates. While complex multi-site mutant prediction is typically handled by specialized models (e.g., ECNET [29],ProteinNPT [30] and PRIME[31]), these models could offer multi-site mutant predictions based on training on single-site mutants. Venus-Maxwell is thus positioned as a crucial tool to provide high-quality, early-stage single-site predictions, which can then be used for experimental validation and as input for downstream multi-mutant prediction models, thereby accelerating the entire directed evolution pipeline.

Besides, we have observed that existing mutation ΔΔ*G* datasets often exhibit significant redundancy between training and test sets, as well as misalignment between mutant and wild-type sequences. To address these issues, we constructed a high-quality dataset derived from public resources, which underwent rigorous de-duplication and redundancy reduction based on sequence alignment. As a result, we obtained strictly separated training and test sets, containing over 226K and 12K curated mutation entries, respectively, ensuring a fair evaluation of generalization. The difference between this dataset and those in previous studies [13] lies in an intensive manual verification process to validate the sequence-mutation consistency.

Utilizing this meticulously constructed ΔΔ*G* dataset for evaluation, we demonstrate that fine-tuning with our Venus-Maxwell framework significantly enhances the predictive capabilities of various PLMs. For example, when using ESM-IF as the base model, the fine-tuned ESM-IF not only outperforms the sequence-to-label paradigm (which uses ESM-IF to generate features per mutant for regression) on the ΔΔ*G* prediction task but also surpasses ThermoMPNN, a current state-of-the-art model designed explicitly for ΔΔ*G* prediction, while achieving over 10 times faster inference.

To summarize, our main contributions are as follows:

### A matrix-based sequence-to-landscape framework

Venus-Maxwell offers a new efficient and broadly applicable approach for fine-tuning diverse PLMs on ΔΔ*G* prediction task. This approach uniquely maximizes the utilization of pre-trained evolutionary knowledge to enable superior stability assessment with minimal computational overhead.

### A qualified mutation ΔΔ*G* dataset

We provide a cleaned protein mutation ΔΔ*G* dataset. The sequences within the training and test sets have been processed to remove duplicates and reduce redundancy, ensuring that their sequence identity remains below 30%.

## 2 Related Work

### Protein language models (PLMs)

Similar to advances in natural language processing, PLMs have evolved into two dominant architectural paradigms: masked language models and auto-regressive models. Masked language models, such as ESM series [22, 24, 25, 32, 33], employ bidirectional attention to predict masked residues, capturing rich contextual representations of protein sequences. Auto-regressive models, like ProGen2 [34] and Tranception [35], predict amino acids sequentially, excelling in generative tasks. These pre-trained models have demonstrated success across various downstream applications, particularly in mutation effect prediction and protein design. Through fine-tuning, PLMs can effectively leverage their encoded evolutionary information to enhance protein fitness prediction, supporting directed evolution [29, 31, 36, 37]. Moreover, PLMs often exhibit zero-shot capabilities [22, 24], but this ability is non-directional and not specifically tailored to stability prediction. Our work offers an efficient way to convert this non-directional prediction into a directional one by fine-tuning on ΔΔ*G* mutation datasets.

### Physics-based methods for ΔΔ*G* prediction

Early efforts in protein stability prediction relied on physics-based methods or statistical analysis [7, 8, 9]. Rosetta [7], a widely adopted framework, employs Cartesian space sampling methods to predict stability changes based on high-resolution structural data. Similarly, FoldX [8] estimates ΔΔ*G* of mutants using an empirically derived energy function calibrated against experimental mutagenesis data. Such methods are often highly interpretable but are limited by computational complexity and input data precision, leading to lower accuracy and speed compared to contemporary deep learning methods, as we present in this work.

### Deep learning-based methods for ΔΔ*G* prediction

Prior to the development of pre-trained PLMs, several deep learning models have been designed to reduce the bias in predicting stabilizing mutations [10, 11, 12]. The application of PLM further advanced ΔΔ*G* prediction. For example, Stability Oracle [13] integrates a pretrained graph transformer model with a data augmentation approach based on thermodynamic permutations to address the challenges of data bias and model generalization. Leveraging the knowledge overlap between sequence recovery and stability optimization tasks, ThermoMPNN [14] utilizes the embeddings from ProteinMPNN [38] for transfer learning, resulting in a fast and robust ΔΔ*G*prediction. Despite their success, these models often require complex architectures and extensive computational resources. Our approach, however, offers a more general and efficient means of facilitating stability prediction with enhanced speed and accuracy, consequently improving its applicability to a range of directed evolution contexts.

## 3 Method

### 3.1 Preliminary

#### Protein Language Model (PLM)

The PLM, whether it is a GPT-style generative model or a BERT-style de-noising model, can be viewed as a parameterized function *f*_*θ*_ that maps a protein sequence **A** = (*a*_1_, *a*_2_, …, *a*_*n*_) to a probability distribution matrix **P** ∈ ℝ^*n*×|*A*|^:

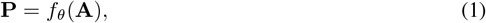

where *n* is the sequence length and |*A*| is the size of the amino acid alphabet (typically 20 for standard amino acids). The element in the *i*-th row and *j*-th column of **P** is the probability that the *i*-th amino acid in the protein sequence is the *j*-th amino acid in the alphabet. Besides, PLMs can be extended by incorporating protein structural condition **S**, modifying the Equation 1 to 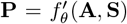. For simplicity, we omit the structural condition in the following, as it does not alter the framework.

#### PLM for mutation scoring

PLMs are typically trained on large-scale protein databases, primarily composed of naturally occurring proteins, which exhibit higher stability due to evolutionary selection. This enables PLMs to predict mutation effects using a “zero-shot” approach by comparing the probabilities of wild-type and mutant amino acids [24]. For a mutation at position *t* from amino acid *a*_*t*_ to *b*_*t*_, the predicted zero-shot score is:

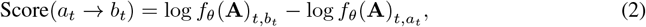

where a higher score indicates a positive mutant. However, the score is often *undirected* (it could reflect activity, stability, or other properties).Notably, for inverse folding models, this specific log-likelihood ratio has been mathematically derived to possess a free energy interpretation directly related to ΔΔ*G* [39]. This provides a strong theoretical foundation for its use. We therefore fine-tune the PLM using ΔΔ*G* data to enable it to explicitly predict mutation ΔΔ*G*.

### 3.2 ΔΔ*G* Landscape Construction

The ΔΔ*G* landscape of a given protein is a real-valued matrix **Y** ∈ **R**^*n*×|A|^, where **Y**_*i,j*_ is the ΔΔ*G* value resulting from mutating the amino acid at position *i* to the *j*-th amino acid. However, in practice, it is usually infeasible to experimentally measure ΔΔ*G* for all possible mutants, leading to a sparse landscape. To track which entries have been experimentally measured, we define a binary mask matrix **M** ∈ *{*0, 1*}*^*n*×|*A*|^, where **M**_*i,j*_ = 1 if **Y**_*i,j*_ is available, and **M**_*i,j*_ = 0 otherwise.

### 3.3 Framework Architecture

The overview of Venus-Maxwell is shown in Figure 2. Specifically, by reformulating zero-shot mutation scoring (Equation 2) as matrix-driven scoring, Venus-Maxwell enables PLMs to compute the mutation landscape in a single forward pass. Crucially, the process is differentiable, allowing PLMs to be fine-tuned on protein mutation ΔΔ*G* landscapes through backpropagation.

**Figure 2.**
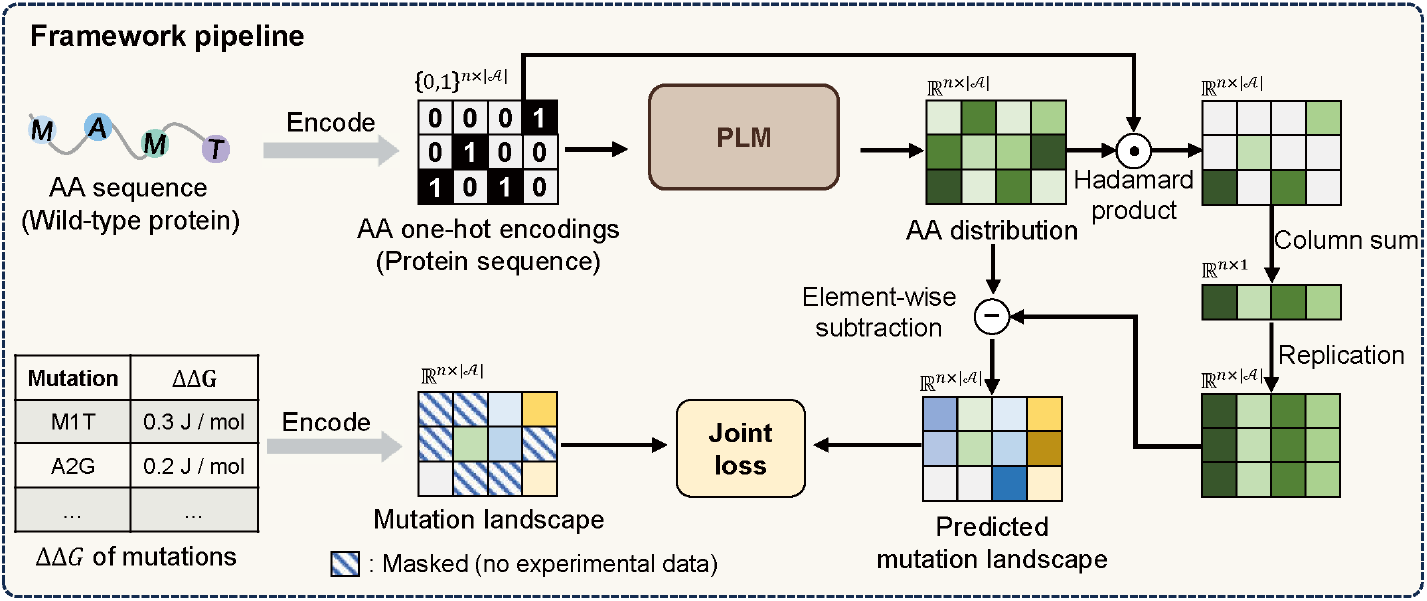
Overview of Venus-Maxwell. Venus-Maxwell utilizes matrix-driven scoring to enable PLMs to learn the mutation landscape of a protein through a single forward-backward computation process.

For a protein sequence **A** = (*a*_1_, *a*_2_, …, *a*_*n*_), Venus-Maxwell computes the landscape as follows:

#### Sequence encoding

The sequence **A** is first tokenized into a one-hot encoding matrix **X** ∈ {0, 1} ^*n*×|A|^. For each position *i*, the row **X**_*i*_ is a one-hot vector where the entry corresponding to the wild-type amino acid *a*_*i*_ is set to 1, and all other entries are set to 0.

#### Probability prediction

The PLM computes a log-probability matrix **Q** ∈ ℝ^*n*×|*A*|^:

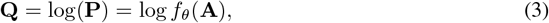

where *f*_*θ*_ is the parameterized PLM function, and the logarithm is applied element-wise. Each entry **Q**_*i,j*_ represents the log-probability of amino acid *a*_*j*_ at position *i*.

#### Wild-type probability extraction

To isolate the log-probabilities associated with the wild-type amino acids, we compute the Hadamard product (element-wise multiplication) between the log-probability matrix **Q** and the one-hot encoding matrix **X**, yielding a matrix **Q**_wt_ ∈ **R**^*n*×|*A*|^:

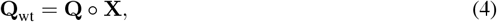

where ○ is the Hadamard product. Since **X**_***i***,***j***_ = 1 only for the wild-type amino acid *a*_*i*_ at position *i*, **Q**_wt_ retains log-probabilities at wild-type positions and zeros elsewhere.

#### Wild-type vector construction

The matrix **Q**_wt_ is then aggregated to produce a compact representation of the wild-type probabilities by summing **Q**_wt_ over the amino acid dimension:

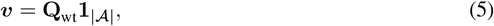

where **1**_|*A*|_ ∈ ℝ^|*A*|^ is a unity vector and *v*_*i*_ is the log-probability of the wild-type amino acid *a*_*i*_.

#### Wild-type matrix replication

To enable comparison between wild-type and mutant probabilities, the wild-type log-probability vector ***v*** is replicated along the column dimension |*A*| times to form a matrix 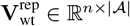. Specifically, we form 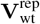 through multiplication of ***v*** with the unity vector 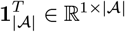:

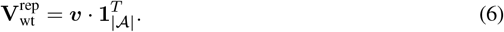

In this matrix, each row *i* contains the log-probability of the wild-type amino acid *a*_*i*_, repeated across all columns *j*.

#### Mutation landscape computation

Finally, we subtract the replicated wild-type matrix 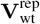 from the log-probability matrix **Q** to obtain the mutation landscape matrix **L** ∈ ℝ^*n*×|*A*|^:

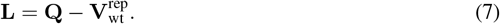

Each entry **L**_*i,j*_ represents the log-probability difference between the *j*-th amino acid and the wild-type amino acid at position *i*. To formalize this computation, we define a function *g*_*θ*_ as follows:

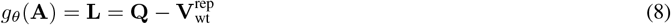

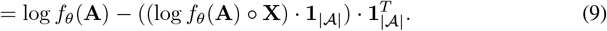

It is clear that *g*_*θ*_ is differentiable because the protein language model *f*_*θ*_ is differentiable. Additionally, the mutation score is equal to the zero-shot scoring formula shown in Equation 2. We also provide the algorithm and pseudo-code of Venus-Maxwell for training and landscape prediction in Appendix A.1 and Appendix A.2.

#### 3.3.1 Training objective

The training objective ensures the predicted ΔΔ*G* landscape closely aligns with the ground-truth ΔΔ*G* landscape. Given a protein sequence **X**, the experimentally measured mutation stability landscape **Y** ∈ ℝ^*n*×|*A*|^, and a binary mask **M** ∈ {0, 1} ^*n*×|*A*|^ indicating observed entries, the model predicts a landscape **L** = *g*_*θ*_(**A**).

The training loss comprises two components: Ranking loss ℒ_Ranking_ and mean squared error loss ℒ_MSE_. The **first** component encourages correct ranking of mutations by minimizing the negative Pearson correlation between the predicted and measured ΔΔ*G* values:

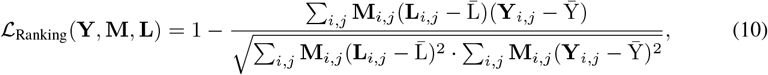

where 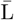 and 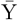 represent the mean predicted and ground-truth ΔΔ*G* values over the masked positions. The **second** component, a mean squared error (MSE) loss, aligns absolute values:

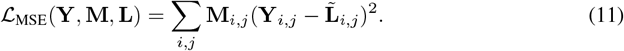

Here, 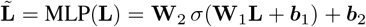 is the output of a two-layer multilayer perceptron (MLP), where *σ*(·) denotes SELU activation function [40]. This transformation adjusts for the scale mismatch between log-probability scores and experimental ΔΔ*G* values. The parameters **W**_1_, **W**_2_, ***b***_1_, and ***b***_2_ are learnable. Specifically, the learnable weights have dimensions **W**_1_ ∈ ℝ^*V* ×*D*^ and **W**_2_ ∈ ℝ^*D*×1^, where *V* is the PLM’s vocabulary size and *D* is the hidden dimension.

The total loss is a weighted sum of both components:

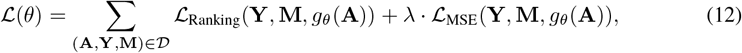

where *λ* is a hyperparameter weighting factor that balances the two loss components. *D* is the training dataset, comprising multiple protein sequences annotated with a sparse mutation landscape and a corresponding binary mask. Each data point in *D* is a tuple (**A, Y, M**). The training objective is to minimize the total loss: *θ*^*\**^ = arg min_*θ*_ ℒ(*θ*). Since *g*_*θ*_ is differential, we can utilize backpropagation and gradient descent to optimize the parameters *θ*, which are initialized from the pre-trained model weights. The specific optimization details, hyperparameter selection, and implementation settings are described in Section 5.1 and Appendix A.3.

## 4 Datasets Building

We collected a large protein mutation ΔΔ*G* dataset, which includes 573 protein sequences and 239,408 mutations (average 418 mutations per protein) to evaluate the performance of Venus-Maxwell. All data have been manually verified to ensure consistency between mutant and wild-type sequences, and duplicates were removed to maintain data integrity.

### Test set

The test set is a compilation of mutation ΔΔ*G* data that we are currently able to collect, including P53 [17], Myoglobin [17], SSym [17], S669 [41], S8754 [42], M1261 [42], vb1432 [43], Fireprotdb [44] and Thermomutdb [45]. Duplicates within the datasets are addressed by selecting the ΔΔ*G* measurement with the highest absolute value. This was intended to address the neutral bias and move the ΔΔ*G* predictions from a model away from 0. After removing duplicates, the test set contains 12,443 mutations across 308 proteins, namely 308 sparse mutation ΔΔ*G* landscape. This dataset, named Test12K, is organized by protein to enable per-protein prediction and evaluation, consistent with benchmarks like ProteinGym [27]. To ensure sufficient data for meaningful per-protein analysis, only proteins with at least five mutations were included.

### Training Set

The training set is derived from a large-scale dataset containing 272K protein mutation sequences, denoted as cDNA272K [46]. This dataset is a commonly used training dataset in the task of protein stability prediction [13]. To prevent potential data leakage issues, we used the MMSeqs2 tool [47] to filter the training set, as illustrated in Figure 3 (A). Specifically, we removed all sequences from the training set that share a sequence identity of 30% or higher with any sequence in the test set. This threshold 30% is chosen to maintain a significant diversity of sequences, as it lies within the “twilight zone”, a region where 95% of protein pairs have distinct structural folds [48]. The resulting training set, Train226K, consists of 226K sequences with less than 30% sequence similarity to any sequences in the test set. All datasets are stored in a protein-specific format to facilitate model training and evaluation, and all sequences were folded using ColabFold 2.5 [49] to generate structural input for PLMs if needed.

**Figure 3.**
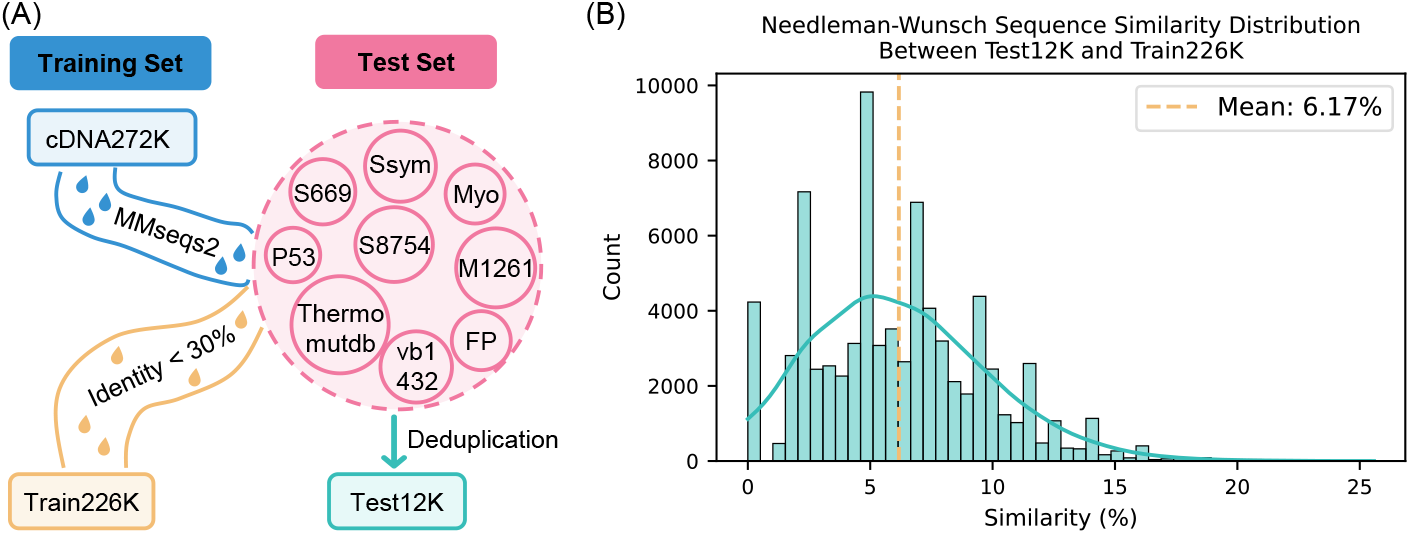
Dataset details. *(A):* Dataset construction process of Train226K and Test12K. *(B):* Global sequence alignment using Needleman-Wunsch confirmed the low sequence similarity between Test12K and Train226K.

To further validate the separation between training and test sets, we performed pairwise global sequence alignment using the Needleman-Wunsch algorithm [50]. As shown in Figure 3 (B), the similarity distribution has a mean of 6.17%, with over 88% of test-training pairs exhibiting less than 10% similarity. This low overlap confirms that our deduplication effectively reduces data leakage risks and enhances the model’s ability to generalize to unseen sequences.

## 5 Experiments and Results

### 5.1 Experiments Setup

The ability of Venus-Maxwell to improve protein stability prediction (ΔΔ*G*) was systematically evaluated across a wide range of PLMs. We strategically selected representative PLMs from two distinct categories: (1) structure-aware PLMs (ESM-IF and ProSST), which incorporate structural information either explicitly or through hybrid architectures, with ESM-IF demonstrating superior zero-shot performance in stability prediction tasks [22] and ProSST representing an emerging paradigm that efficiently combines sequence and structural information [23]; and (2) sequence-only PLMs (ESM-1b, ESM2, ESM-1v), which are widely adopted in protein engineering applications and serve as pure sequence-based model representatives [24, 51, 52, 33, 53].

#### Hyper-parameter settings

For all Venus-Maxwell enhanced models, we utilized the Adam optimizer [54]. All key hyperparameters were selected through a rigorous 5-fold cross-validation procedure on the training set (Train226K) to prevent test set leakage. Based on this analysis, we set the initial learning rate to 5 × 10^−5^, the loss weighting factor *λ* to 0.1, and the MLP hidden dimension *D* to *V* (the PLM’s vocabulary size). The optimal number of training epochs (approximately 7, determined via early stopping with a patience of 5 epochs) was also identified during this crossvalidation process. The final models were then trained on the complete Train226K dataset for the determined optimal number of epochs. A detailed description of the cross-validation methodology, hyperparameter search spaces, and full ablation results for each choice is provided in Appendix A.3. All experiments were conducted on a PC with an NVIDIA RTX 4090 GPU.

#### Evaluation metrics

To rigorously assess model performance, we employed per-protein evaluation followed by averaging across the dataset. For each protein, we calculated Pearson (*ρ*_*p*_) and Spearman (*ρ*_*s*_) correlation coefficients between predicted and experimental ΔΔ*G*values, capturing linear and monotonic relationships, respectively. For classification of stabilizing vs. destabilizing mutations (threshold at ΔΔ*G* = 0), we computed AUC-ROC and F1 scores. This per-protein averaging approach prevents bias toward proteins with more mutations, ensuring balanced assessment across diverse protein structures [27, 28].

#### Baselines

We compared Venus-Maxwell against: (1) ThermoMPNN, the current state-of-the-art neural network model [55], used with its publicly available checkpoint^2^. Our training set is a subset of the training set of ThermoMPNN, so there are no data leakage issues. (2)FoldX and Rosetta, two representative physics-based computational methods; and (3) Embedding transfer baselines using ProSST and ESM-IF, denoted as “ProSST + MLP” and “ESM-IF + MLP”. The hyperparameter settings of baseline models are shown in Appendix A.4.

### 5.2 Model Performance

We evaluated Venus-Maxwell’s performance on the Test12K dataset (Section 4), comprising 12,443 mutations across 308 diverse proteins. As shown in Figure 4 (A), Venus-Maxwell enhances zeroshot performance across multiple PLMs, achieving an average Spearman correlation improvement of 0.143 (Wilcoxon signed-rank test, *p <* 0.01). With ESM-IF, Venus-Maxwell reaches a Spearman correlation of 0.517, followed by ProSST at 0.479, highlighting the importance of structure-aware representations in protein stability prediction tasks [27, 28]. This improvement stems from matrix-driven scoring (Equation 8), which maps sequences to mutation landscapes in one forward pass.

**Figure 4.**
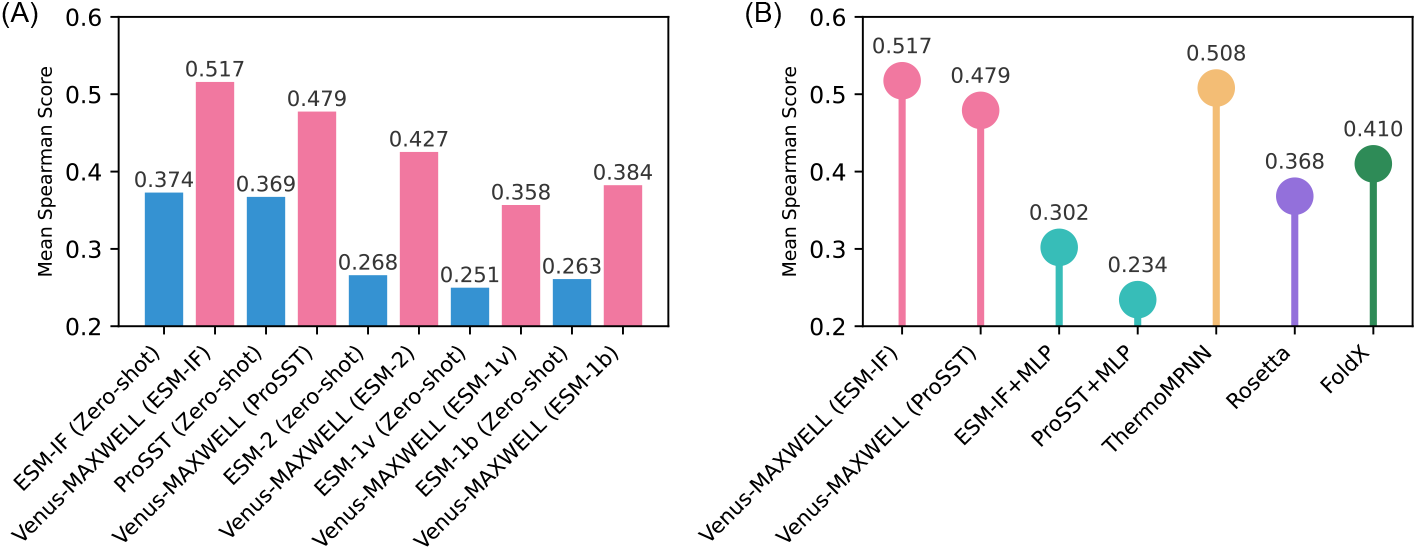
Performance comparison of Venus-Maxwell. *(A):* Comparison of Spearman correlation scores between zero-shot predictions and Venus-Maxwell enhanced predictions across different PLMs. The improvement from zero-shot to Venus-Maxwell is statistically significant (Wilcoxon signed-rank test, *p <* 0.01). *(B):* Comparison with existing mutation prediction methods. Venus-Maxwell (ESM-IF) outperforms ThermoMPNN (Wilcoxon signed-rank test, *p <* 0.01). All evaluations were performed on the Test12K dataset, with per-protein correlations averaged across the entire set.

When compared to existing methods (Figure 4 (B)), embedding transfer methods struggle under the low similarity between Train226K and Test12K (Spearman: 0.302/0.234). The physics-based Rosetta and FoldX method also demonstrates moderate performance. In contrast, the specialized architecture of ThermoMPNN delivers strong results (Spearman: 0.508). Notably, Venus-Maxwell with ESM-IF surpasses ThermoMPNN (Spearman: 0.517, *p <* 0.01) while Venus-Maxwell with ProSST achieves comparable performance, without requiring architectural modifications to the base PLMs. To further validate these findings on established public benchmarks, we also benchmarked performance on specific, well-known subsets within Test12K (e.g., p53, s669, Myoglobin) against ThermoMPNN and Stability Oracle. As detailed in Appendix A.5, Venus-Maxwell consistently outperforms both SOTA models on these individual datasets, confirming its robust and state-of-the-art performance. These results demonstrate that Venus-Maxwell ‘s matrix-based landscape approach effectively captures global stability patterns, outperforming both traditional regression methods and physics-based approaches by leveraging evolution pattern knowledge from pre-trained PLMs. For more details of comparison on additional metrics, please refer to Table A4 in Appendix A.5.

Furthermore, we investigated whether the inherent sparsity of the training landscapes adversely affects generalization. We conducted an ablation study by progressively removing random portions of the training data. The results show that Venus-Maxwell’s performance is exceptionally robust, maintaining a superior performance (Spearman: 0.501) even when 90% of the training mutants are removed. This demonstrates that the model learns generalizable stability principles from the diverse collection of proteins, rather than overfitting to sparse local landscapes. The full details of this sparsity analysis are also presented in Appendix A.5.

To ensure that the superior performance of Venus-Maxwell with ESM-IF and ProSST, both of which utilize structural information, does not stem from structural data leakage, we analyzed structural similarity between Train226K and Test12K using TM-align [56]. The average TM-score between training and test set structures is approximately 0.3, indicating low structural similarity. Furthermore, the weak correlation between prediction accuracy and structural similarity (see more details in Figure A1 and Appendix A.6) confirms Venus-Maxwell’s robust generalization ability across diverse protein structures.

### 5.3 Prediction Speed Analysis

We evaluated Venus-Maxwell’s inference efficiency in the context of directed evolution, where rapid screening of single-site mutants is crucial. Testing on single-site saturation mutagenesis of 10 random proteins from Test12K (averaged over five sampling runs), Venus-Maxwell demonstrates remarkable speed improvements over baseline methods.

Despite maintaining full model capacity (142 million parameters for Venus-Maxwell (ESM-IF) and 119 million parameters for Venus-Maxwell (ProSST)),Venus-Maxwell achieves exceptional inference efficiency. As shown in Table 1, Venus-Maxwell(ESM-IF) predicts 20,050 mutants per second, while Venus-Maxwell(ProSST) reaches 85,471 mutants per second, outperforming ThermoMPNN (2,088 mutants/s) by over 10-fold and MLP-based baselines (9–51 mutants/s) by orders of magnitude. The efficiency improvement remained robust across sequence lengths (100–1000 residues), as detailed in Table A7 and A8 in Appendix A.7, where we further analyze inference speed variations with sequence length, underscoring Venus-Maxwell’s capability for parallel prediction of mutation landscapes. Notably, the training efficiency is similarly improved: Venus-Maxwell (ESM-IF) completes one training epoch on Train226K in 41 seconds, over 200 times faster than embedding transfer methods, despite fine-tuning the entire PLM (see Figure A2 in Appendix A.8 for details). These efficiency gains, combined with the predictive accuracy reported in Section 5.2, highlight Venus-Maxwell’s scalability for high-throughput directed evolution.

**Table 1:** Efficiency comparison of stability prediction methods.

| Model | Trainable parameters | Prediction time (s) | Mutants per second |
| --- | --- | --- | --- |
| Venus-MAXWELL (ESM-IF) | 142 M | $2.11 \pm 0.04$ | $20050 \pm 714$ |
| Venus-MAXWELL (ProSST) | 119 M | $0.53 \pm 0.02$ | $85471 \pm 3512$ |
| ESM-IF + MLP | 132 K | $4744.76 \pm 335.9$ | $9 \pm 1$ |
| ProSST + MLP | 296 K | $807.77 \pm 7.53$ | $51 \pm 1$ |
| ThermoMPNN | 2.68 M | $20.27 \pm 1.81$ | $2088 \pm 7$ |

### 5.4 Ablation Study

To dissect the contribution of Venus-Maxwell’s components, we conducted ablation studies using ESM-IF as the base model, evaluating performance on Test12K (Table 2).Our analysis reveals:

**Table 2:** Ablation studies of Venus-Maxwell (ESM-IF) on Test12K datasets.

| Model | $\bar{\rho}_s$ | $\bar{\rho}_p$ | $\bar{F1}$ | $\bar{AUC}$ |
| --- | --- | --- | --- | --- |
| Venus-MAXWELL (ESM-IF) | **0.517** | **0.542** | **0.427** | **0.678** |
| Venus-MAXWELL (ESM-IF) (No $\mathcal{L}_{MSE}$) | 0.506 | 0.528 | 0.330 | 0.638 |
| Venus-MAXWELL (ESM-IF) (No $\mathcal{L}_{Ranking}$) | 0.496 | 0.518 | 0.410 | 0.671 |
| Venus-MAXWELL (ESM-IF) (ESMFold) | 0.492 | 0.519 | 0.389 | 0.662 |
| Venus-MAXWELL (ESM-IF without pretrain) | 0.202 | 0.234 | 0.382 | 0.501 |
| ESM-IF + MLP | 0.302 | 0.331 | 0.383 | 0.627 |
| Zero-shot | 0.375 | 0.376 | 0.218 | 0.575 |
| Zero-shot (ESMFold) | 0.354 | 0.360 | 0.217 | 0.574 |
| ThermoMPNN | 0.508 | 0.529 | 0.371 | 0.646 |

#### Ablation on training objective

Removing the MSE module (“No ℒ_MSE_”) maintained strong mutation ranking (Spearman: 0.506) but decreased classification performance (F1: 0.330, AUC: 0.638). Conversely, removing the ranking loss component (“No ℒ_Ranking_”) better preserved classification metrics (F1: 0.410, AUC: 0.671) while showing reduced correlation scores (Spearman: 0.496). Notably, both variants remained competitive with ThermoMPNN, demonstrating the robustness of our matrix-based approach even with partial architecture.

#### Ablation on structure quality

Using ESMFold-generated structures instead of ColabFold slightly reduced performance [51]. As a result, if the structure is predicted using ESMFold instead of ColabFold, both our model (Venus-Maxwell (ESMFold)) and the zero-shot method experienced a slight decrease in scoring accuracy. Specifically, the Spearman correlation dropped from 0.517 to 0.492 for our model and from 0.375 to 0.354 for the zero-shot method, respectively.

#### Ablation on pre-training impact

Random initialization of ESM-IF severely degraded performance (Spearman: 0.202), demonstrating the crucial role of pre-trained weights in capturing evolutionary patterns. All models with pre-trained weights outperformed both embedding transfer (Spearman: 0.302) and zero-shot baselines (0.375/0.354 for ESM-IF/ESMFold), confirming the effectiveness of our matrix-based landscape approach.

## 6 Conclusion and Limitations

In this work, we introduced Venus-Maxwell, a universal and efficient framework that enhances protein stability prediction while preserving PLMs’ inherent capabilities. Through elegant matrix transformations, our approach enables rapid modeling of the complete mutation landscape, demonstrating superior performance in both ranking and classification metrics. Extensive evaluation demonstrates a computational speedup of three orders of magnitude, establishing a scalable paradigm for protein engineering and stability optimization.

While Venus-Maxwell excels in stability prediction, its matrix-based approach holds some potential for broader applications. For example, the current matrix-based implementation is inherently limited to PLMs using single amino acids as vocabulary tokens, excluding structure-aware tokenization schemes (*e*.*g*., SaProt [57]) or BPE tokenizer [58] (*e*.*g*., ProGPT [59]). Moreover, the framework focuses on single-site saturation mutations now, and extending it to model combinatorial mutation effects poses a key challenge. Future work could address these limitations by developing adaptive tokenization strategies, incorporating multi-site mutation modeling, and evaluating Venus-Maxwell on diverse protein function prediction tasks.

## Acknowledgments and Disclosure of Funding

This work was supported by the National Key Research and Development Program of China (2024YFA0917603), Computational Biology Key Program of Shanghai Science and Technology Commission (23JS1400600), and Science and Technology Innovation Key R&D Program of Chongqing (CSTB2022TIAD-STX0017).

## A Appendix

### A.1 Algorithm

Algorithm 1 outlines the training procedure for Venus-Maxwell. The process iterates through the dataset, computing the predicted mutation landscape *L*_*i*_ from the PLM’s log-probabilities for each protein *A*_*i*_. The model parameters *θ* are then updated using gradient descent based on a combined loss function that incorporates both ranking (ℒ_Ranking_) and absolute value (ℒ_MSE_) objectives.

#### Algorithm 1

Venus-Maxwell for mutation ΔΔ*G* landscape learning

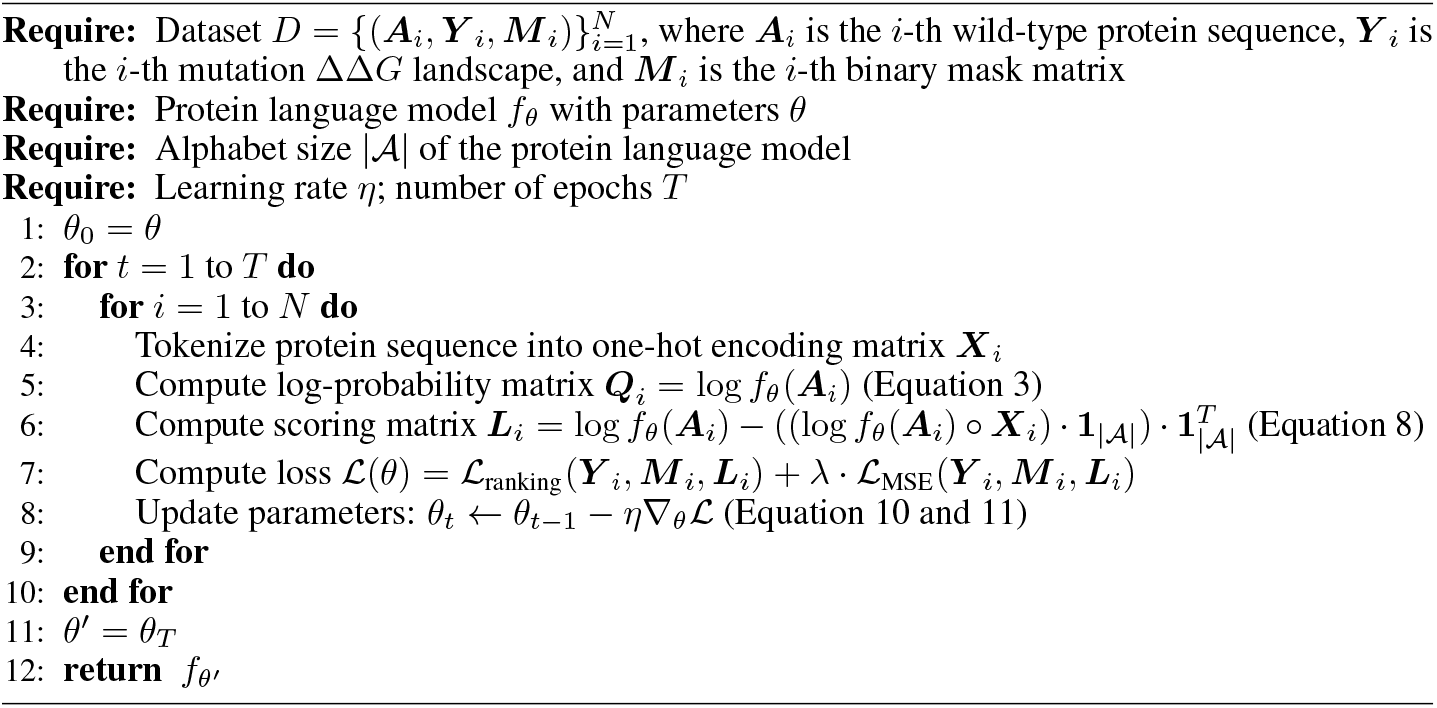

Algorithm 2 details the prediction process using a PLM fine-tuned by Venus-Maxwell. Given a protein sequence *A*, the model performs a single forward pass to compute the log-probability matrix and subsequently derives the mutation ΔΔ*G* landscape *L* based on Equation 8.

#### Algorithm 2

Venus-Maxwell for mutation ΔΔ*G* landscape prediction

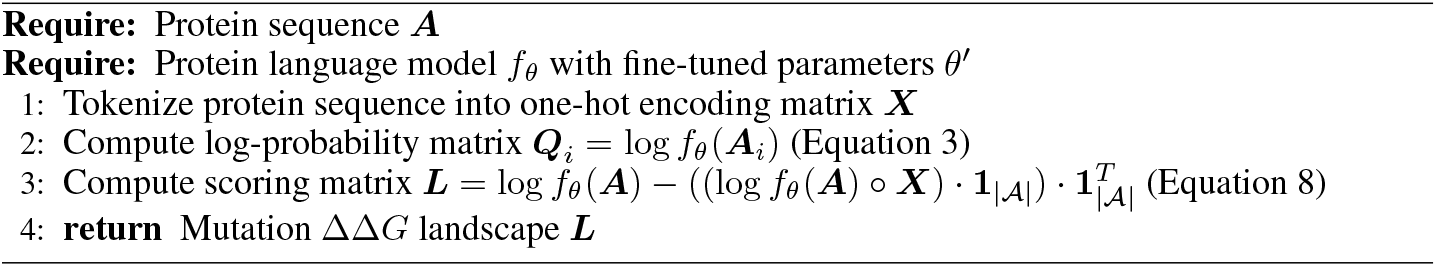

### A.2 Pseudo Code

Algorithm 3 provides a Pytorch-like pseudocode implementation of the core training loop for Venus-Maxwell. It illustrates how input sequences are processed, logits are computed, and transformed into the mutation landscape matrix, and how the ranking and MSE losses are calculated and combined for backpropagation and parameter updates.

#### Algorithm 3

Pseudocode of Venus-Maxwell training in a pytorch-like style.

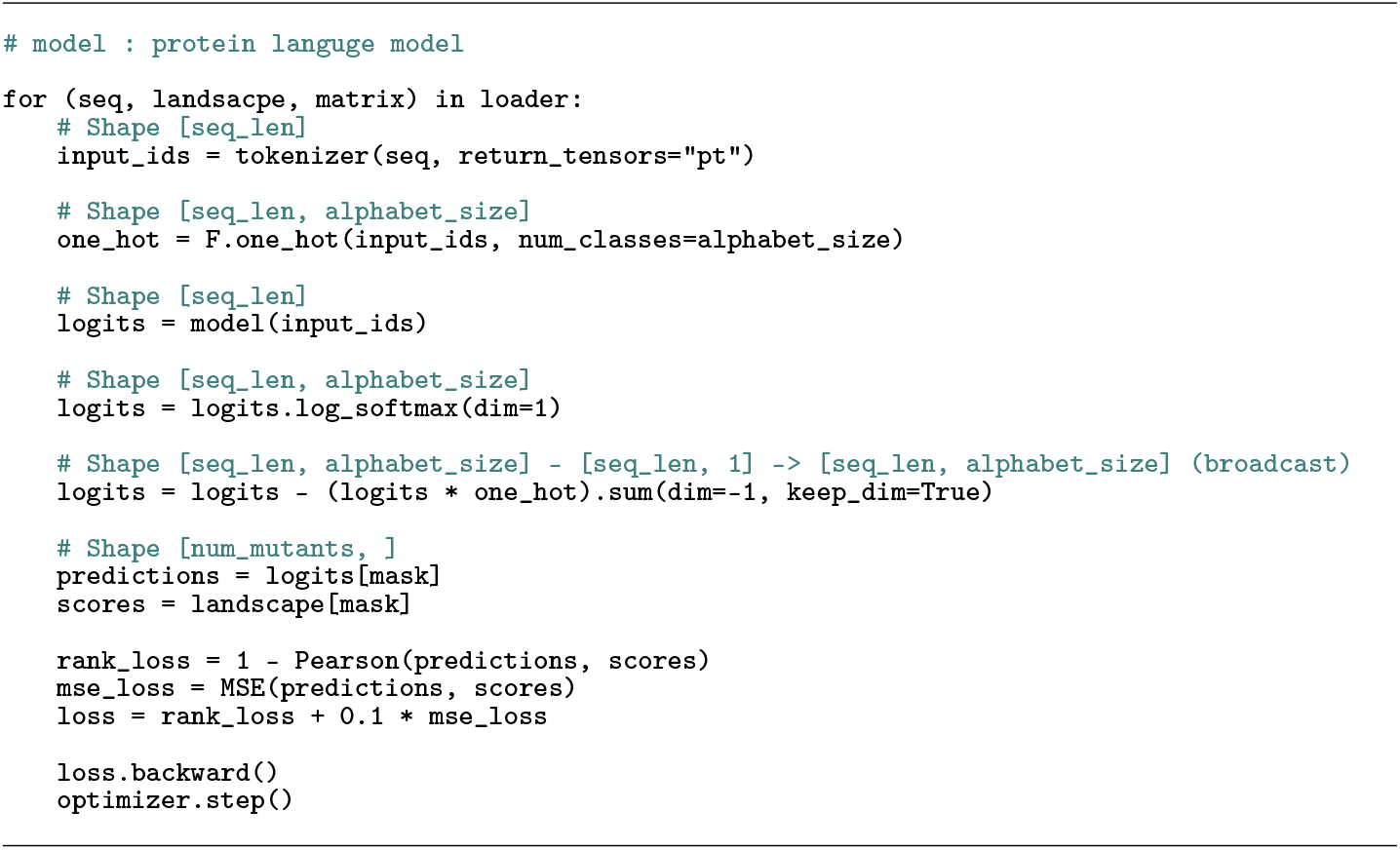

### A.3 Hyperparameter Selection and Ablation Studies

To ensure a fair and robust selection of hyperparameters without test set leakage, we performed 5-fold cross-validation (CV) on the **training dataset**. The training set was split into five folds {*T*_1_, *T*_2_, *T*_3_, *T*_4_, *T*_5_}. For each set of hyperparameters (e.g., a specific learning rate), we followed this procedure:

- For *i* = 1 to 5:
  ‐ Set the validation fold: *D*_val_ = *T*_*i*_.
  ‐ Set the internal training set: *D*_train_internal_ = *D*_train_ \*T*_*i*_.
  ‐ Train the model on *D*_train_internal_ for a fixed number of epochs (e.g., 10 epochs for the initial LR search).
  ‐ Record the best Spearman correlation *ρ* achieved on the validation fold *D*_val_.
- Calculate the average Spearman *ρ* across all 5 folds.

We define the “optimal performance” as this 5-fold mean per-protein Spearman correlation 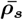. A higher average score indicates a better hyperparameter choice. After identifying the optimal hyperparameters (e.g., learning rate 5 × 10^−5^), we determined the average optimal training epochs (approximately 7) from the CV folds. Finally, we trained a single model on the **entire** training dataset for 7 epochs and reported its performance on the held-out test set (Test12K). The test set was used exclusively for this final evaluation.

#### Ablation for Loss Weighting Factor *λ*

In our loss function (defined in Equation 12), *λ* balances the ranking loss (ℒ_Ranking_) and the MSE loss (ℒ_MSE_). We conducted a sensitivity analysis by training Venus-Maxwell (ESM-IF) with different values of *λ*.

As shown in Table A1, the model achieved the best performance on the test dataset when *λ* = 0.1. This suggests the framework performs optimally when the ranking loss serves as the primary training objective, while the MSE loss acts as a beneficial auxiliary objective to align the scale of the predictions. Based on this analysis, we set *λ* = 0.1 for all other experiments.

#### Ablation for Learning Rate (lr)

**Table A1:** Ablation study on the weighting factor *λ*. We report the average Spearman 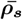 and Pearson 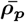 correlation coefficients of Venus-Maxwell (ESM-IF) on the Test12K dataset.

| Weighting Factor ($\lambda$) | $\bar{\rho}_s$ | $\bar{\rho}_p$ |
| --- | --- | --- |
| 0.0 | 0.506 | 0.528 |
| **0.1** | **0.517** | **0.542** |
| 0.2 | 0.514 | 0.540 |
| 0.4 | 0.504 | 0.538 |
| 0.8 | 0.494 | 0.523 |
| 1.0 | 0.486 | 0.519 |

Following the 5-fold CV methodology, we performed a grid search for the optimal learning rate (lr). The results are shown in Table A2. We selected 5 × 10^−5^ as the optimal learning rate, as it demonstrated the best average performance across the validation sets.

**Table A2:**
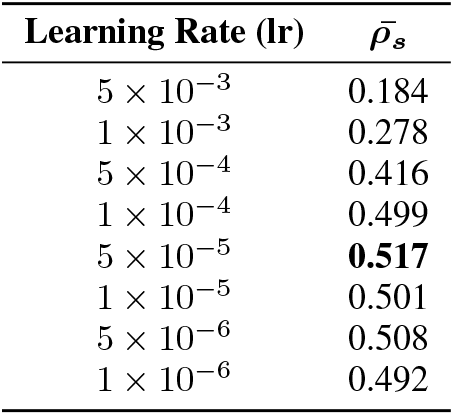
Performance of Venus-Maxwell (ESM-IF) with different learning rates during 5-fold cross-validation. “Avg Spearman *ρ*” refers to the mean Spearman correlation across the 5 validation folds. Best performance is in **bold**.

#### Ablation for MLP Hidden Layer Dimension

The MLP (used in Equation 11) is a two-layer network where the learnable weights have dimensions **W**_1_ ∈ ℝ^*V* ×*D*^ and **W**_2_ ∈ ℝ ^*D*×1^. *V* is the PLM vocabulary size, and *D* is a tunable hidden dimension. We investigated the model’s sensitivity to *D*, using ESM-IF (*V* = 35) as the base model.

As shown in Table A3, the results indicate that the model achieves its best performance when the hidden dimension *D* is set to *V* (35). While performance is relatively stable for *D* up to 512, this analysis validates our choice of *D* = *V*, showing that increasing the MLP’s complexity does not yield further benefits.

**Table A3:** Performance of Venus-Maxwell (ESM-IF) with varying MLP hidden layer dimensions (*D*).

| Hidden Dimension ($D$) | $\bar{\rho}_s$ |
| --- | --- |
| **V (35)** | **0.517** |
| 64 | 0.506 |
| 128 | 0.516 |
| 256 | 0.515 |
| 512 | 0.505 |
| 1024 | 0.487 |

#### Evaluation Metrics and Embedding Transfer Baseline Models Configuration

For embedding transfer, features from frozen PLMs were fed into a three-layer MLP with SELU activations and 0.1 dropout, trained with Adam optimizer (learning rate 1 × 10^−3^, batch size 32). Following the similar training protocol as Venus-Maxwell, the optimal number of training epochs was determined through five-fold cross-validation on the training set, with early stopping during validation (patience of 5 epochs based on Pearson correlation).

### A.5 Model Performance Details

**Table A4:** Model performance across different metrics.

| Model Type | Model | $\bar{\rho}_s$ | $\bar{\rho}_p$ | $\bar{F1}$ | $\bar{AUC}$ |
| --- | --- | --- | --- | --- | --- |
| PLM zero-shot | ESM-IF (Zero-shot) | 0.375 | 0.376 | 0.217 | 0.577 |
|  | ProSST (Zero-shot) | 0.369 | 0.370 | 0.372 | 0.560 |
|  | ESM-1b (Zero-shot) | 0.263 | 0.278 | 0.281 | 0.524 |
|  | ESM-1v (Zero-shot) | 0.251 | 0.256 | 0.295 | 0.527 |
|  | ESM-2 (Zero-shot) | 0.268 | 0.284 | 0.289 | 0.533 |
| Baseline models | ThermoMPNN | 0.508 | 0.529 | 0.371 | 0.647 |
|  | Rosetta | 0.368 | 0.239 | 0.346 | 0.503 |
|  | FoldX | 0.410 | 0.431 | 0.322 | 0.581 |
| Embedding transfer | ESM-IF + MLP | 0.302 | 0.331 | 0.383 | 0.628 |
|  | ProSST + MLP | 0.234 | 0.263 | 0.360 | 0.608 |
| Venus-MAXWELL (PLM) | Venus-MAXWELL (ESM-IF) | **0.517** | **0.542** | **0.427** | **0.678** |
|  | Venus-MAXWELL (ProSST) | 0.479 | 0.502 | 0.405 | 0.604 |
|  | Venus-MAXWELL (ESM-1b) | 0.384 | 0.402 | 0.334 | 0.578 |
|  | Venus-MAXWELL (ESM-1v) | 0.358 | 0.371 | 0.324 | 0.555 |
|  | Venus-MAXWELL (ESM-2) | 0.427 | 0.450 | 0.366 | 0.545 |

Table A4 provides a detailed breakdown of the performance for all evaluated models across the four primary metrics: Mean Spearman correlation 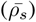, Mean Pearson correlation 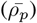, Mean F1 score 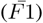, and Mean Area Under the Curve 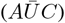. Models are categorized by type (PLM zero-shot, baseline SOTA model, and physical-based method, embedding transfer, and Venus-Maxwell) to facilitate comparison within and across different approaches. The best performance for each metric is highlighted in bold. This comprehensive table allows for a granular assessment of each model’s strengths and weaknesses on the Test12K dataset.

In addition to this comprehensive performance comparison, we further validated our model on specific, established public benchmarks. We clarify that Test12K is a curated meta-benchmark composed of these benchmarks (e.g., p53, s669). To directly address concerns about evaluation solely on our constructed dataset, Table A5 presents a detailed comparison against SOTA models on several individual subsets.

The results show that Venus-Maxwell consistently outperforms both ThermoMPNN and Stability Oracle on these well-known datasets. We deliberately excluded the popular ProteinGym benchmark, as its significant overlap with the training data of both our model and ThermoMPNN would lead to data leakage and invalidate the conclusions.

**Table A5:** Model Performance on Public Benchmarks (Mean per-protein Spearman 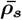).

| Benchmark | Venus-MAXWELL (ESM-IF) | ThermoMPNN | Stability Oracle |
| --- | --- | --- | --- |
| p53 | **0.751** | 0.702 | 0.727 |
| s669 | **0.565** | 0.545 | 0.503 |
| Myoglobin | **0.724** | 0.620 | 0.669 |
| Ssym | **0.602** | 0.516 | 0.567 |

Finally, to address concerns about the inherent sparsity of the training data and its potential to adversely affect generalization, we conducted an ablation study on training set sparsity. We evaluated Venus-Maxwell (ESM-IF) after randomly removing *p* percent of the mutation entries from the Train226K dataset. Performance (Mean per-protein Spearman 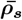) on the Test12K set remains stable even with extreme data removal.

As shown in Table A6, the framework demonstrates exceptional robustness. Performance remains remarkably stable, with a Spearman’s correlation of 0.501 even when using only 10% (23K mutants) of the training data. A significant performance drop only occurs at the extreme of removing 95% of the data. This analysis confirms that our model learns generalizable stability principles from the diverse collection of proteins, rather than overfitting to sparse local landscapes.

**Table A6:** Effect of training set sparcsity on model performance.

| Data Removed ($p$) | Training Mutants | $\bar{\rho}_s$ | $\bar{\rho}_p$ | $\bar{F1}$ | $\bar{AUC}$ |
| --- | --- | --- | --- | --- | --- |
| 0% | 226K | 0.517 | 0.542 | 0.427 | 0.678 |
| 20% | 181K | 0.517 | 0.541 | 0.423 | 0.673 |
| 40% | 136K | 0.515 | 0.541 | 0.421 | 0.675 |
| 60% | 90K | 0.512 | 0.540 | 0.416 | 0.669 |
| 80% | 45K | 0.509 | 0.537 | 0.408 | 0.667 |
| 90% | 23K | 0.501 | 0.523 | 0.402 | 0.660 |
| 95% | 11K | 0.485 | 0.506 | 0.388 | 0.647 |

### A.6 Generalization Ability Analysis Across Structual identity

In constructing the training and test sets, we ensured low sequence similarity via MMseqs2 deduplication and Needleman-Wunsch validation (See Section 4). Since the top-performing Venus-Maxwell enhanced models, ESM-IF and ProSST, leverage protein structure information, we further assessed the structural similarity between the sets to evaluate the models’ extrapolation capability to structurally novel proteins. We employed TM-align [56] to compute TM-scores for each protein in Test12K against all structures in Train226K, using the maximum score as the structural similarity metric.

As illustrated in Figure A1 (A), the maximum TM-scores of the test proteins displayed a tightly clustered distribution, with the overall probability density peaking around a mean value of approximately 0.31. According to established conventions in structural biology, TM-scores below 0.5 indicate distinct folding topologies [60]. This confirms the structural diversity between Train226K and Test12K, establishing a rigorous benchmark for extrapolation.

Further analysis of Venus-Maxwell (ESM-IF) performance relative to structural similarity revealed only a weak correlation (*ρ*_*p*_ = 0.208, as shown in Figure A1 (B)). This finding indicates that the performance enhancement of structure-aware protein language models within the Venus-Maxwell framework is independent of structural similarity to training samples, demonstrating strong generalization capabilities to unseen protein architectures.

**Figure A1:**
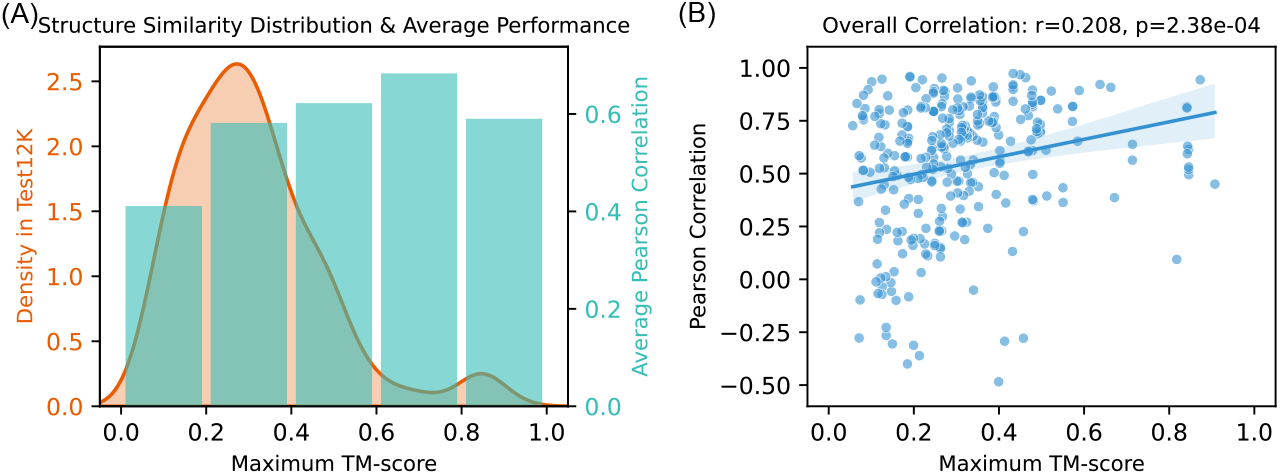
Model generalization of Venus-Maxwell (ESM-IF) against structural dissimilarity. *(A):* Distribution of maximum TM-scores for Test12K proteins relative to Train226K, overlaid with average Pearson correlation per TM-score bin. *(B):*Relationship between maximum TM-score and prediction performance, showing minimal dependence on structural similarity to training proteins.

### A.7 Prediction Speed Scaling with Sequence Length

**Table A7:** Prediction time (s) across sequence length ranges for single-site mutagenesis.

| Model | 0-200 | 200-400 | 400-600 | 600+ |
| --- | --- | --- | --- | --- |
| Venus-MAXWELL (ProSST) | 0.30±0.10 | 0.37±0.13 | 0.27±0.02 | 0.30±0.07 |
| Venus-MAXWELL (ESM-IF) | 1.33±0.10 | 1.65±0.10 | 1.78±0.13 | 2.19±0.16 |
| ThermoMPNN | 4.41±0.04 | 13.64±0.01 | 20.64±0.11 | 30.89±0.13 |
| ProSST + MLP | 162.19±0.87 | 539.72±0.67 | 844.52±3.68 | 1347.73±6.83 |
| ESM-IF + MLP | 421.44±1.63 | 2846.02±1.13 | 6255.54±13.25 | 13554.11±7.27 |

**Table A8:** Mutants per second across different sequence length ranges for single-site mutagenesis.

| Model | 0-200 | 200-400 | 400-600 | 600+ |
| --- | --- | --- | --- | --- |
| Venus-MAXWELL (ProSST) | 30451±8141 | 82336±27736 | 159005±8935 | 217882±43378 |
| Venus-MAXWELL (ESM-IF) | 6513±501 | 16887±973 | 23993±1705 | 29425±2155 |
| ThermoMPNN | 1957±17 | 2038±2 | 2058±11 | 2080±9 |
| ProSST + MLP | 53.22±0.29 | 51.51±0.06 | 50.29±0.22 | 47.67±0.24 |
| ESM-IF + MLP | 20.48±0.08 | 9.77±0.01 | 6.79±0.01 | 4.74±0.00 |

To assess the scalability of Venus-Maxwell’s inference efficiency across diverse protein sizes, we evaluated its prediction speed for single-site saturation mutagenesis on proteins with different sequence length ranges from the Test12K dataset. For each sequence length range (0–200, 200–400, 400–600, and 600+ residues), we randomly selected five proteins, performed single-site saturation mutagenesis, and measured inference speed across all models. This process was repeated five times to compute the mean and standard deviation of prediction times and throughput.

As shown in Table A7, prediction times for all methods except Venus-Maxwell (ProSST) increase with sequence length. For instance, ThermoMPNN’s prediction time rises from 4.41 ± 0.04 s (0–200 residues) to 30.89 ± 0.13 s (600+ residues), and MLP-based baselines (ProSST-MLP and ESM-IF-MLP) exhibit even steeper increases, reaching up to 13,554.11 ± 7.27 s for ESM-IF-MLP on sequences exceeding 600 residues. This trend is driven by two factors: (1) longer sequences result in a higher total number of single-site mutants, and (2) Transformer-based PLMs require significantly more time to encode longer sequences due to their quadratic complexity with respect to sequence length.

In contrast, Venus-Maxwell (ProSST) maintains rather stable prediction times (0.27–0.37 s across all ranges), likely due to its remarkable high throughput, which remains well above the demands of current sequence lengths and mutation counts. Table A8 further validates this observation. The throughput of embedding transfer methods decreases with sequence length—from 53.22 ± 0.29 to 47.67 ± 0.24 mutants/s for ProSST + MLP and from 20.48 ± 0.08 to 4.74 ± 0.00 mutants/s for ESM-IF + MLP—reflecting their need to re-encode each mutant sequence, making them highly sensitive to sequence length. ThermoMPNN, leveraging ProteinMPNN’s structure to reduce encoding overhead, achieves a relatively stable throughput of approximately 2,000 mutants/s (1,957–2,080 mutants/s), but this appears to approach its practical upper limit.

Conversely, both Venus-Maxwell (ProSST) and Venus-Maxwell (ESM-IF) exhibit significant throughput increases with sequence length, reaching orders-of-magnitude higher performance. This scalability is likely attributable to Venus-Maxwell’s optimized matrix processing architecture (see Section 3), which efficiently handles the whole mutation landscapes.

### A.8 Model Training Speed

This Figure A2 highlights the training efficiency of Venus-Maxwell compared with other methods, using ESM-IF and ProSST as base models. Embedding transfer methods, which require encoding each mutant sequence during forward propagation, are significantly constrained in training speed. For ESM-IF and ProSST, processing 100,000 mutant sequences takes over 2,000 seconds, with a single training epoch exceeding one hour. ThermoMPNN substantially reduces training time by encoding only the wild-type sequence, processing 100,000 mutations in under 300 seconds. While the training efficiency of Venus-Maxwellis further 20 times faster than ThermoMPNN, completing an epoch on the entire Train225K dataset in approximately 40 seconds.

**Figure A2:**
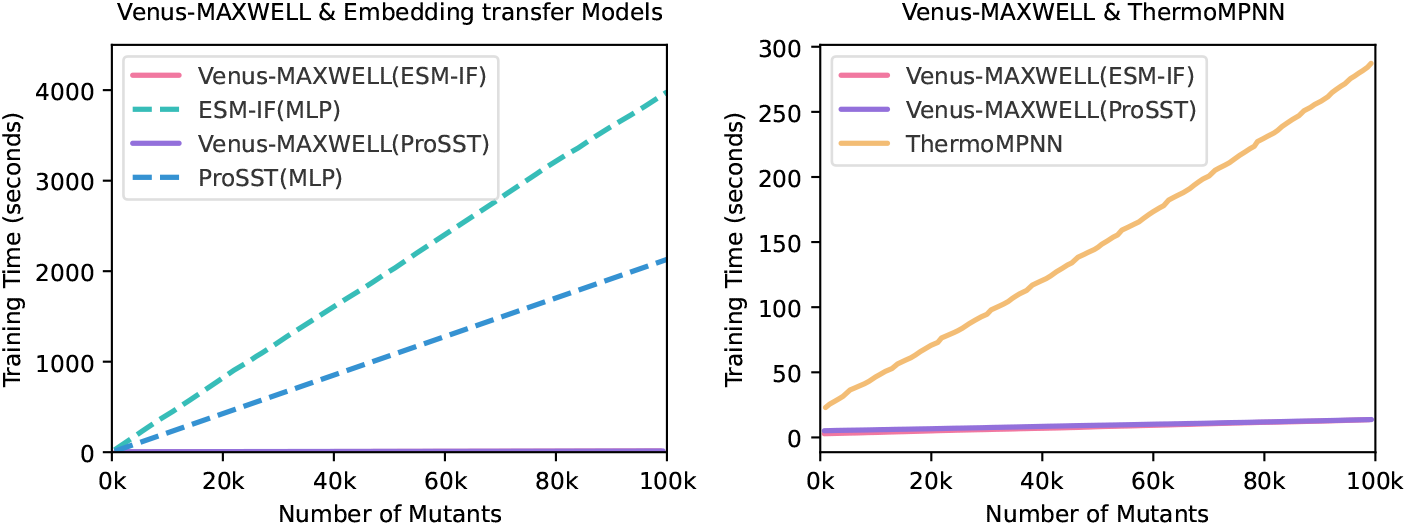
Training time analysis. *(A):* Comparison between Venus-Maxwell models and their embedding transfer (MLP) counterparts for both ESM-IF and ProSST architectures. *(B):* Performance of Maxwell implementations contrasted with ThermoMPNN

## Footnotes

^2^https://github.com/Kuhlman-Lab/ThermoMPNN

## References

[1] Sasha B Ebrahimi and Devleena Samanta. Engineering protein-based therapeutics through structural and chemical design. Nature Communications, 14(1):2411, 2023.

[2] Steven M Jay and Richard T Lee. Protein engineering for cardiovascular therapeutics: untapped potential for cardiac repair. Circulation research, 113(7):933–943, 2013.

[3] Michaela Gebauer and Arne Skerra. Engineered protein scaffolds as next-generation therapeutics. Annual review of pharmacology and toxicology, 60(1):391–415, 2020.

[4] Elizabeth L Bell, William Finnigan, Scott P France, Anthony P Green, Martin A Hayes, Lorna J Hepworth, Sarah L Lovelock, Haruka Niikura, Sílvia Osuna, Elvira Romero, et al. Biocatalysis. Nature Reviews Methods Primers, 1(1):1–21, 2021.

[5] Zhoutong Sun, Qian Liu, Ge Qu, Yan Feng, and Manfred T Reetz. Utility of b-factors in protein science: interpreting rigidity, flexibility, and internal motion and engineering thermostability. Chemical reviews, 119(3):1626–1665, 2019.

[6] Fan Jiang, Jiahao Bian, Hao Liu, Song Li, Xue Bai, Lirong Zheng, Sha Jin, Zhuo Liu, Guang-Yu Yang, and Liang Hong. Creatinase: Using increased entropy to improve the activity and thermostability. The Journal of Physical Chemistry B, 127(12):2671–2682, 2023.

[7] Rebecca F Alford, Andrew Leaver-Fay, Jeliazko R Jeliazkov, Matthew J O’Meara, Frank P DiMaio, Hahnbeom Park, Maxim V Shapovalov, P Douglas Renfrew, Vikram K Mulligan, Kalli Kappel, et al. The rosetta all-atom energy function for macromolecular modeling and design. Journal of chemical theory and computation, 13(6):3031–3048, 2017.

[8] Joost Schymkowitz, Jesper Borg, Francois Stricher, Robby Nys, Frederic Rousseau, and Luis Serrano. The foldx web server: an online force field. Nucleic acids research, 33(suppl_2):W382–W388, 2005.

[9] Shuangye Yin, Feng Ding, and Nikolay V Dokholyan. Eris: an automated estimator of protein stability. Nature methods, 4(6):466–467, 2007.

[10] Bian Li, Yucheng T Yang, John A Capra, and Mark B Gerstein. Predicting changes in protein thermodynamic stability upon point mutation with deep 3d convolutional neural networks. PLoS computational biology, 16(11):e1008291, 2020.

[11] Huali Cao, Jingxue Wang, Liping He, Yifei Qi, and John Z Zhang. Deepddg: predicting the stability change of protein point mutations using neural networks. Journal of chemical information and modeling, 59(4):1508–1514, 2019.

[12] Lijun Quan, Qiang Lv, and Yang Zhang. Strum: structure-based prediction of protein stability changes upon single-point mutation. Bioinformatics, 32(19):2936–2946, 2016.

[13] Daniel J Diaz, Chengyue Gong, Jeffrey Ouyang-Zhang, James M Loy, Jordan Wells, David Yang, Andrew D Ellington, Alexandros G Dimakis, and Adam R Klivans. Stability oracle: a structure-based graph-transformer framework for identifying stabilizing mutations. Nature Communications, 15(1):6170, 2024.

[14] Henry Dieckhaus, Michael Brocidiacono, Nicholas Z Randolph, and Brian Kuhlman. Transfer learning to leverage larger datasets for improved prediction of protein stability changes. Proceedings of the National Academy of Sciences, 121(6):e2314853121, 2024.

[15] John Jumper, Richard Evans, Alexander Pritzel, Tim Green, Michael Figurnov, Olaf Ronneberger, Kathryn Tunyasuvunakool, Russ Bates, Augustin Žídek, Anna Potapenko, et al. Highly accurate protein structure prediction with alphafold. nature, 596(7873):583–589, 2021.

[16] Ziyi Zhou, Liang Zhang, Yuanxi Yu, Banghao Wu, Mingchen Li, Liang Hong, and Pan Tan. Enhancing efficiency of protein language models with minimal wet-lab data through few-shot learning. Nature Communications, 15(1):5566, 2024.

[17] Bian Li, Yucheng T Yang, John A Capra, and Mark B Gerstein. Predicting changes in protein thermodynamic stability upon point mutation with deep 3d convolutional neural networks. PLoS computational biology, 16(11):e1008291, 2020.

[18] Lasse M Blaabjerg, Maher M Kassem, Lydia L Good, Nicolas Jonsson, Matteo Cagiada, Kristoffer E Johansson, Wouter Boomsma, Amelie Stein, and Kresten Lindorff-Larsen. Rapid protein stability prediction using deep learning representations. Elife, 12:e82593, 2023.

[19] Shuyu Wang, Hongzhou Tang, Peng Shan, Zhaoxia Wu, and Lei Zuo. Pros-gnn: predicting effects of mutations on protein stability using graph neural networks. Computational Biology and Chemistry, 107:107952, 2023.

[20] Yuanxi Yu, Fan Jiang, Bozitao Zhong, Liang Hong, and Mingchen Li. Entropy-driven zero-shot deep learning model selection for viral proteins. Physical Review Research, 7(1):013229, 2025.

[21] Liang Zhang, Kuan Luo, Ziyi Zhou, Yuanxi Yu, Fan Jiang, Banghao Wu, Mingchen Li, and Liang Hong. A deep retrieval-enhanced meta-learning framework for enzyme optimum ph prediction. Journal of Chemical Information and Modeling, 65(7):3761–3770, 2025.

[22] Chloe Hsu, Robert Verkuil, Jason Liu, Zeming Lin, Brian Hie, Tom Sercu, Adam Lerer, and Alexander Rives. Learning inverse folding from millions of predicted structures. In International conference on machine learning, pages 8946–8970. PMLR, 2022.

[23] Mingchen Li, Yang Tan, Xinzhu Ma, Bozitao Zhong, Huiqun Yu, Ziyi Zhou, Wanli Ouyang, Bingxin Zhou, Pan Tan, and Liang Hong. ProSST: Protein language modeling with quantized structure and disentangled attention. In The Thirty-eighth Annual Conference on Neural Information Processing Systems, 2024.

[24] Joshua Meier, Roshan Rao, Robert Verkuil, Jason Liu, Tom Sercu, and Alex Rives. Language models enable zero-shot prediction of the effects of mutations on protein function. Advances in neural information processing systems, 34:29287–29303, 2021.

[25] Zeming Lin, Halil Akin, Roshan Rao, Brian Hie, Zhongkai Zhu, Wenting Lu, Nikita Smetanin, Robert Verkuil, Ori Kabeli, Yaniv Shmueli, et al. Evolutionary-scale prediction of atomic-level protein structure with a language model. Science, 379(6637):1123–1130, 2023.

[26] Ahmed Elnaggar, Michael Heinzinger, Christian Dallago, Ghalia Rehawi, Yu Wang, Llion Jones, Tom Gibbs, Tamas Feher, Christoph Angerer, Martin Steinegger, et al. Prottrans: Toward understanding the language of life through self-supervised learning. IEEE transactions on pattern analysis and machine intelligence, 44(10):7112–7127, 2021.

[27] Pascal Notin, Aaron Kollasch, Daniel Ritter, Lood Van Niekerk, Steffanie Paul, Han Spinner, Nathan Rollins, Ada Shaw, Rose Orenbuch, Ruben Weitzman, et al. Proteingym: Large-scale benchmarks for protein fitness prediction and design. Advances in Neural Information Processing Systems, 36:64331–64379, 2023.

[28] Liang Zhang, Hua Pang, Chenghao Zhang, Song Li, Yang Tan, Fan Jiang, Mingchen Li, Yuanxi Yu, Ziyi Zhou, Banghao Wu, et al. Venusmuthub: A systematic evaluation of protein mutation effect predictors on small-scale experimental data. Acta Pharmaceutica Sinica B, 2025.

[29] Yunan Luo, Guangde Jiang, Tianhao Yu, Yang Liu, Lam Vo, Hantian Ding, Yufeng Su, Wesley Wei Qian, Huimin Zhao, and Jian Peng. Ecnet is an evolutionary context-integrated deep learning framework for protein engineering. Nature communications, 12(1):5743, 2021.

[30] Pascal Notin, Ruben Weitzman, Debora Marks, and Yarin Gal. Proteinnpt: Improving protein property prediction and design with non-parametric transformers. Advances in Neural Information Processing Systems, 36:33529–33563, 2023.

[31] Fan Jiang, Mingchen Li, Jiajun Dong, Yuanxi Yu, Xinyu Sun, Banghao Wu, Jin Huang, Liqi Kang, Yufeng Pei, Liang Zhang, et al. A general temperature-guided language model to design proteins of enhanced stability and activity. Science Advances, 10(48):eadr2641, 2024.

[32] Thomas Hayes, Roshan Rao, Halil Akin, Nicholas J Sofroniew, Deniz Oktay, Zeming Lin, Robert Verkuil, Vincent Q Tran, Jonathan Deaton, Marius Wiggert, et al. Simulating 500 million years of evolution with a language model. Science, page eads0018, 2025.

[33] Alexander Rives, Joshua Meier, Tom Sercu, Siddharth Goyal, Zeming Lin, Jason Liu, Demi Guo, Myle Ott, C Lawrence Zitnick, Jerry Ma, et al. Biological structure and function emerge from scaling unsupervised learning to 250 million protein sequences. Proceedings of the National Academy of Sciences, 118(15):e2016239118, 2021.

[34] Erik Nijkamp, Jeffrey A Ruffolo, Eli N Weinstein, Nikhil Naik, and Ali Madani. Progen2: exploring the boundaries of protein language models. Cell systems, 14(11):968–978, 2023.

[35] Pascal Notin, Mafalda Dias, Jonathan Frazer, Javier Marchena-Hurtado, Aidan N Gomez, Debora Marks, and Yarin Gal. Tranception: protein fitness prediction with autoregressive transformers and inference-time retrieval. In International Conference on Machine Learning, pages 16990–17017. PMLR, 2022.

[36] Kaiyi Jiang, Zhaoqing Yan, Matteo Di Bernardo, Samantha R Sgrizzi, Lukas Villiger, Alisan Kayabolen, BJ Kim, Josephine K Carscadden, Masahiro Hiraizumi, Hiroshi Nishimasu, et al. Rapid in silico directed evolution by a protein language model with evolvepro. Science, page eadr6006, 2024.

[37] Mingchen Li, Liqi Kang, Yi Xiong, Yu Guang Wang, Guisheng Fan, Pan Tan, and Liang Hong. Sesnet: sequence-structure feature-integrated deep learning method for data-efficient protein engineering. Journal of Cheminformatics, 15(1):12, 2023.

[38] Justas Dauparas, Ivan Anishchenko, Nathaniel Bennett, Hua Bai, Robert J Ragotte, Lukas F Milles, Basile IM Wicky, Alexis Courbet, Rob J de Haas, Neville Bethel, et al. Robust deep learning–based protein sequence design using proteinmpnn. Science, 378(6615):49–56, 2022.

[39] Jes Frellsen, Maher M Kassem, Tone Bengtsen, Lars Olsen, Kresten Lindorff-Larsen, Jesper Ferkinghoff-Borg, and Wouter Boomsma. Zero-shot protein stability prediction by inverse folding models: a free energy interpretation. arXiv preprint arXiv:2506.05596, 2025.

[40] Günter Klambauer, Thomas Unterthiner, Andreas Mayr, and Sepp Hochreiter. Self-normalizing neural networks. Advances in neural information processing systems, 30, 2017.

[41] Corrado Pancotti, Silvia Benevenuta, Giovanni Birolo, Virginia Alberini, Valeria Repetto, Tiziana Sanavia, Emidio Capriotti, and Piero Fariselli. Predicting protein stability changes upon single-point mutation: a thorough comparison of the available tools on a new dataset. Briefings in Bioinformatics, 23(2):bbab555, 2022.

[42] Yunxin Xu, D. Liu, and Haipeng Gong. Improving the prediction of protein stability changes upon mutations by geometric learning and a pre-training strategy. Nature Computational Science, pages 1–11, 2024.

[43] Yang Yang, Siddhaling Urolagin, Abhishek Niroula, Xuesong Ding, Bairong Shen, and Mauno Vihinen. Pon-tstab: protein variant stability predictor. importance of training data quality. International journal of molecular sciences, 19(4):1009, 2018.

[44] Jan Stourac, Juraj Dubrava, Milos Musil, Jana Horackova, Jiri Damborsky, Stanislav Mazurenko, and David Bednar. Fireprotdb: database of manually curated protein stability data. Nucleic acids research, 49(D1):D319–D324, 2021.

[45] Joicymara S Xavier, Thanh-Binh Nguyen, Malancha Karmarkar, Stephanie Portelli, Pâmela M Rezende, Joao PL Velloso, David B Ascher, and Douglas EV Pires. Thermomutdb: a thermodynamic database for missense mutations. Nucleic acids research, 49(D1):D475–D479, 2021.

[46] Kotaro Tsuboyama, Justas Dauparas, Jonathan Chen, Elodie Laine, Yasser Mohseni Behbahani, Jonathan J Weinstein, Niall M Mangan, Sergey Ovchinnikov, and Gabriel J Rocklin. Mega-scale experimental analysis of protein folding stability in biology and design. Nature, 620(7973):434–444, 2023.

[47] Martin Steinegger and Johannes Söding. Mmseqs2 enables sensitive protein sequence searching for the analysis of massive data sets. Nature biotechnology, 35(11):1026–1028, 2017.

[48] Burkhard Rost. Twilight zone of protein sequence alignments. Protein engineering, 12(2):85–94, 1999.

[49] Milot Mirdita, Konstantin Schütze, Yoshitaka Moriwaki, Lim Heo, Sergey Ovchinnikov, and Martin Steinegger. Colabfold: making protein folding accessible to all. Nature methods, 19(6):679–682, 2022.

[50] Vladimir Likic. The needleman-wunsch algorithm for sequence alignment. Lecture given at the 7th Melbourne Bioinformatics Course, Bi021 Molecular Science and Biotechnology Institute, University of Melbourne, pages 1–46, 2008.

[51] Zeming Lin, Halil Akin, Roshan Rao, Brian Hie, Zhongkai Zhu, Wenting Lu, Allan dos Santos Costa, Maryam Fazel-Zarandi, Tom Sercu, Sal Candido, et al. Language models of protein sequences at the scale of evolution enable accurate structure prediction. BioRxiv, 2022:500902, 2022.

[52] Lin Tang. Large model predicts variant effects. Nature Methods, 20(10):1448–1448, 2023.

[53] Benjamin J Livesey and Joseph A Marsh. Advancing variant effect prediction using protein language models. Nature genetics, 55(9):1426–1427, 2023.

[54] Diederik P Kingma. Adam: A method for stochastic optimization. arXiv preprint arXiv:1412.6980, 2014.

[55] Antoni Beltran, Xiang’er Jiang, Yue Shen, and Ben Lehner. Site-saturation mutagenesis of 500 human protein domains. Nature, pages 1–10, 2025.

[56] Yang Zhang and Jeffrey Skolnick. Tm-align: a protein structure alignment algorithm based on the tm-score. Nucleic acids research, 33(7):2302–2309, 2005.

[57] Jin Su, Chenchen Han, Yuyang Zhou, Junjie Shan, Xibin Zhou, and Fajie Yuan. Saprot: Protein language modeling with structure-aware vocabulary. In The Twelfth International Conference on Learning Representations, 2024.

[58] Philip Gage. A new algorithm for data compression. The C Users Journal, 12(2):23–38, 1994.

[59] Noelia Ferruz, Steffen Schmidt, and Birte Höcker. Protgpt2 is a deep unsupervised language model for protein design. Nature communications, 13(1):4348, 2022.

[60] Jinrui Xu and Yang Zhang. How significant is a protein structure similarity with tm-score= 0.5? Bioinformatics, 26(7):889–895, 2010.

